# Continuous and discrete quantity discrimination in tortoises

**DOI:** 10.1101/425678

**Authors:** Andrea Gazzola, Giorgio Vallortigara, Daniele Pellitteri-Rosa

## Abstract

The ability to estimate quantity, which is crucially important in several aspects of animal behaviour (e.g., foraging), has been extensively investigated in most taxa, with the exception of reptiles. The few studies available, in lizards, report lack of spontaneous discrimination of quantity, which may suggest that reptiles could represent an exception in numerical abilities among vertebrates. We investigated the spontaneous ability of Hermann’s tortoises (*Testudo hermanni*) to select the larger quantity of food items. Tortoises showed able to choose the larger food item when exposed with two options differing in size (0.25, 0.50, 0.67 and 0.75 ratio) and when presented with two groups differing in numerousness (1 versus 4, 2 versus 4, 2 versus 3 and 3 versus 4 items). The tortoises succeeded in both size and numerousness discrimination, and their performance appeared to depend on the ratio of items to be discriminated (thus following Weber’s Law). These findings in chelonians provide evidence of an ancient system for the extrapolation of numerical magnitudes from given sets of elements, shared among vertebrates.

## Introduction

Animals use information which is potentially available in their environment in order to survive, find food and reproduce. An example is the ability to discriminate between sets of physical objects (e.g., food items, conspecifics, predators, refuges) making use of discrete (countable) or continuous quantities. Evidence showed that when animals make non-symbolic quantity judgments, their accuracy is limited by the ratio between the numerical values being compared, as indicated in Weber’s Law (review in Ferrigno and Cantlon, 2017). For example, animals might be identically accurate at choosing the larger of two sets when the numerical choices have the same ratio (e.g., 5 versus 10, 10 versus 20, or 50 versus 100: 1.0 Weber fraction) but show a decrease in accuracy with lower ratios (e.g., 4 versus 5, 16 versus 20, and 40 versus 50: 0.25 Weber fraction).

This ratio-dependent pattern of success and failure has been documented in warm-blooded vertebrates in a number of species (mammals, e.g., Cantlon and Brannon, 2007; Utrata, Virányi, Range, 2012; Vonk, Beran, 2012; birds: e.g., Rugani et al, 2009; Pepperberg, 2006; Ditz and Nieder, 2016; reviews in Vallortigara, 2014; 2017). Less clear is the evidence for quantity discrimination in cold-blooded vertebrates. Amphibians (Uller et al., 2003; Stancher et al., 2015) and fish (Agrillo et al., 2012; Potrich et al., 2015) showed quantity judgements that vary as a function of ratio in accordance with Weber’s law. In reptiles, in contrast, ruin lizards (*Podarcis siculus*) proved able to spontaneously discriminate between the surface area of two food items of different size, but failed when food was presented in sets of discrete items differing in numerousness (Miletto Petrazzini et al., 2017). The lizards showed very poor performance also in experiments involving explicit training: six out of 10 discriminated 1 vs. 4 items; among these, only one was capable of learning a 2 vs. 4 discrimination (Miletto Petrazzini et al., 2018). This is clearly in contrast with evidence collected in fish and amphibians (above) in similar tasks.

In this study, we explored for the first time the numerical competence in chelonians, and particularly in *Testudo hermanni*, in discriminating between quantities differing in number or size. We adopted a similar protocol to that used in the experiment conducted on lizards (Miletto Petrazzini et al., 2017) by presenting tortoises with four different combinations of food items. Each combination represented a choice test between two items and was performed with four proportional differences in magnitude (i.e. ratio s: 0.25, 0.50, 0.67, 0.75), both for number and size (see methods below).

## 2. Material and Methods

### (a) Subjects

We collected 25 adult Hermann’s tortoises, *Testudo hermanni*, from the naturalistic area of “Oasi di Sant’Alessio”, located roughly 20 km south of Milan, in Northern Italy. In order to determine sexual adultness, we measured carapace length of all turtles using a digital calliper (accuracy ± 0.1 mm). Sexual dimorphism is noticeable in this species: males are smaller and their plastron is concave, which allows them to mount females during mating (Willemsen and Hailey, 2003). In our experiments we selected 16 males and 3 females (carapace length mean ± ES, males: 167.7 ± 4.3 mm, females: 200.0 ± 26.2 mm). The total sample consisted of 16 subjects for number discrimination experiment (14 males, 2 females), and 15 subjects in size discrimination experiment (13 males, 2 females). Of these, twelve tortoises (11 males and 1 female) were used in both experiments.

### (b) Experimental apparatus and visual stimulus

The tortoises were tested in an outdoor arena consisting of a tunnel that served both as starting zone and approach area to the stimuli and a wider testing compartment where stimuli were presented during the trials (figure 1). Subjects could observe the stimuli from the starting of the trial and throughout the tunnel before entering the testing compartment. Visual stimuli were represented by *Solanum lycopersicum* (San Marzano tomato variety) slices placed on two supports arranged in the testing compartment. Slices were presented in a symmetrical position compared to the longitudinal axis of the tunnel. Since olfactory cues might represent a signal of attraction (Petrazzini et al., 2017), four tomato slices were included in the experimental compartment and hidden from the sight of the tortoises (see the electronic supplementary material, S1, for additional details).

**Fig 1.**
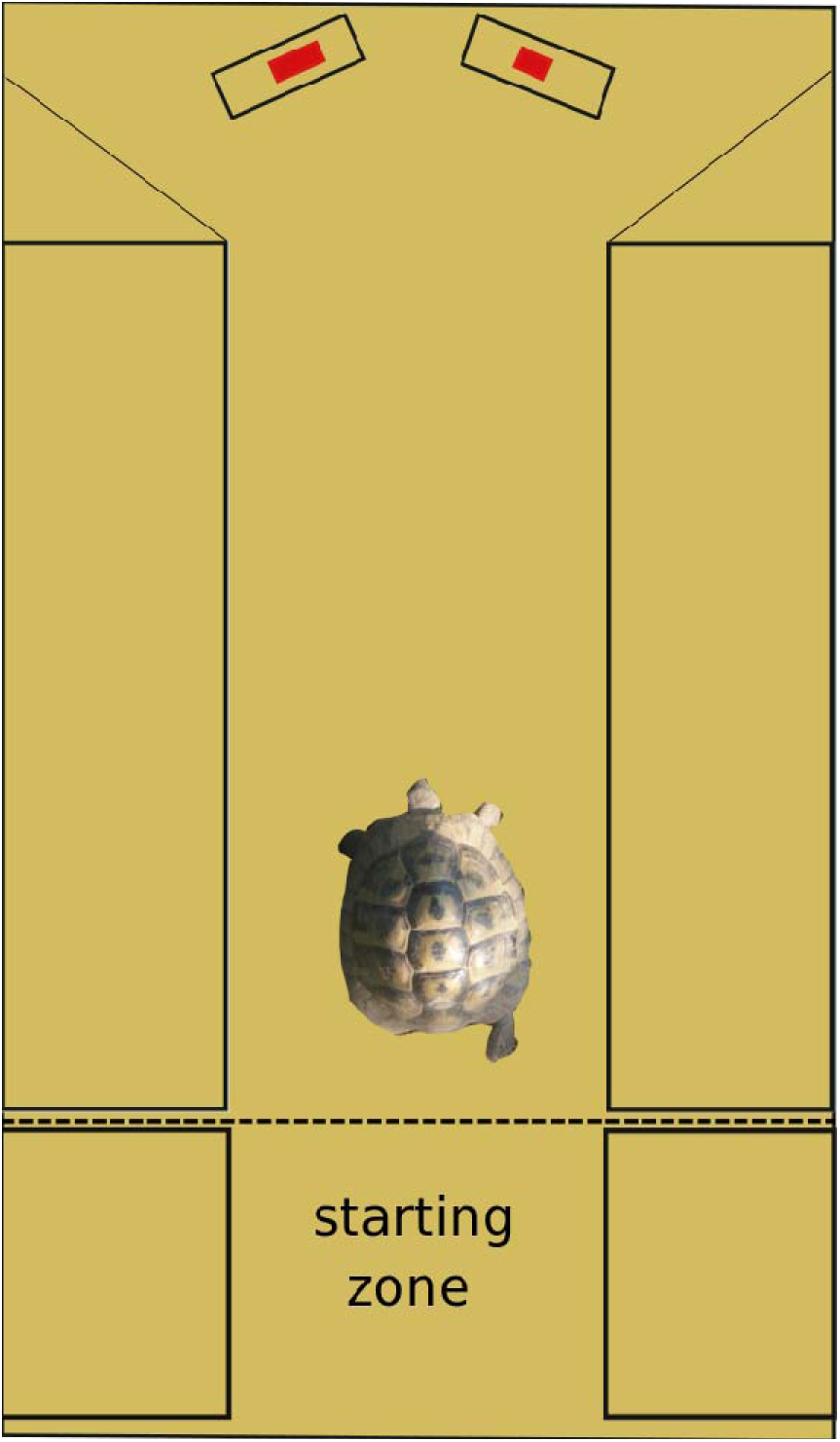
Experimental set up used in the study. For details see electronic supplementary materials.

### (c) Experimental procedures

We applied the same general procedure and the same setup in the two different experiments (number and size experiment; see the electronic supplementary material, S1, for details). A 5-day acclimation period was necessary in order to allow the tortoises to become familiar with the experimental apparatus, and subsequently be tested. Each tortoise was presented with two food items, which were different both in numerousness (number experiment) and in dimension (size experiment). In each testing trial, the subject was allowed a single choice between the two items. We considered the choice made as the first attempt to eat any tomato slice of any group. When the subject approached the slices at the distance of about 1 cm, the choice was considered made. The tortoise was then removed from the arena and placed back into its enclosure. However, in order to avoid any possible learning effect, the tortoises were not allowed to eat the tomatoes (Stancher et al., 2015). In this way, the subject performed the task in the general context of a spontaneous choice. If after 3 min the tortoise had not approached any stimulus, the response was discarded, and the trial was repeated. The left–right position of the larger stimulus was presented in a pseudo-random sequence to exclude the possible effect of lateralization (Rogers et al, 2013). All different food item combinations were presented in a mixed sequence in both of the experiments.

### (d) Number discrimination experiment

We investigated the tortoises’ choices between two groups of food items of the same size (circular tomato slices, diameter = 2 cm) but differing in number. We adopted four numerical comparisons: 1 versus 4, 2 versus 4, 2 versus 3 and 3 versus 4, representing four different ratios within each combination (0.25, 0.50, 0.67 and 0.75, respectively). Each tortoise was tested in a total of 60 trials (15 for each discrimination) over 13 days.

### (e) Size discrimination experiment

In this experiment, we explored the ability of tortoises to distinguish between food items of different size. We observed subjects in their spontaneous preference between combinations (1 versus 1) of differently sized tomato slices (range from 2 to 8 cm^2^), with four size ratios within each combination (0.25, 0.50, 0.67 and 0.75, as in the first experiment). The tortoises underwent a total of 60 trials for each subject (15 for each discrimination) over a period of 13 days.

## Results

Overall, the tortoises showed a significant preference for each combination presented with, both for larger food items and higher numerosity (figure 3). Both the main effects in the mixed model performed on the index of choice (number of choice for larger item / total number of choice) were significant (combination (ratios): *χ*^2^=31.21, df=3, p<0.001; experiment: *χ*^2^=4.87, df=1, p=0.027) but not their interaction (*χ*^2^=2.10, df=3, p=0.55). The pattern analysis, performed including the combination as a numerical variable in the model, showed a main effect of the combination *χ*^2^ = 16.70, p<0.001) and of the experiment (*χ*^2^ = 4.74, p=0.030) but no significant effect of their interaction (*χ*^2^ = 0.69, p=0.40). The analysis conducted separately on the single experiment showed a decline in performance for both number (*β* ± se = −0.03 ± 0.01, t=−2.22, p=0.03) and size experiments (*β* ± se = −0.05 ± 0.01, t=−3.66, p<0.001; figure 2), but their decrease was not different (t=−0.83, p=0.41).

**Fig 2.**
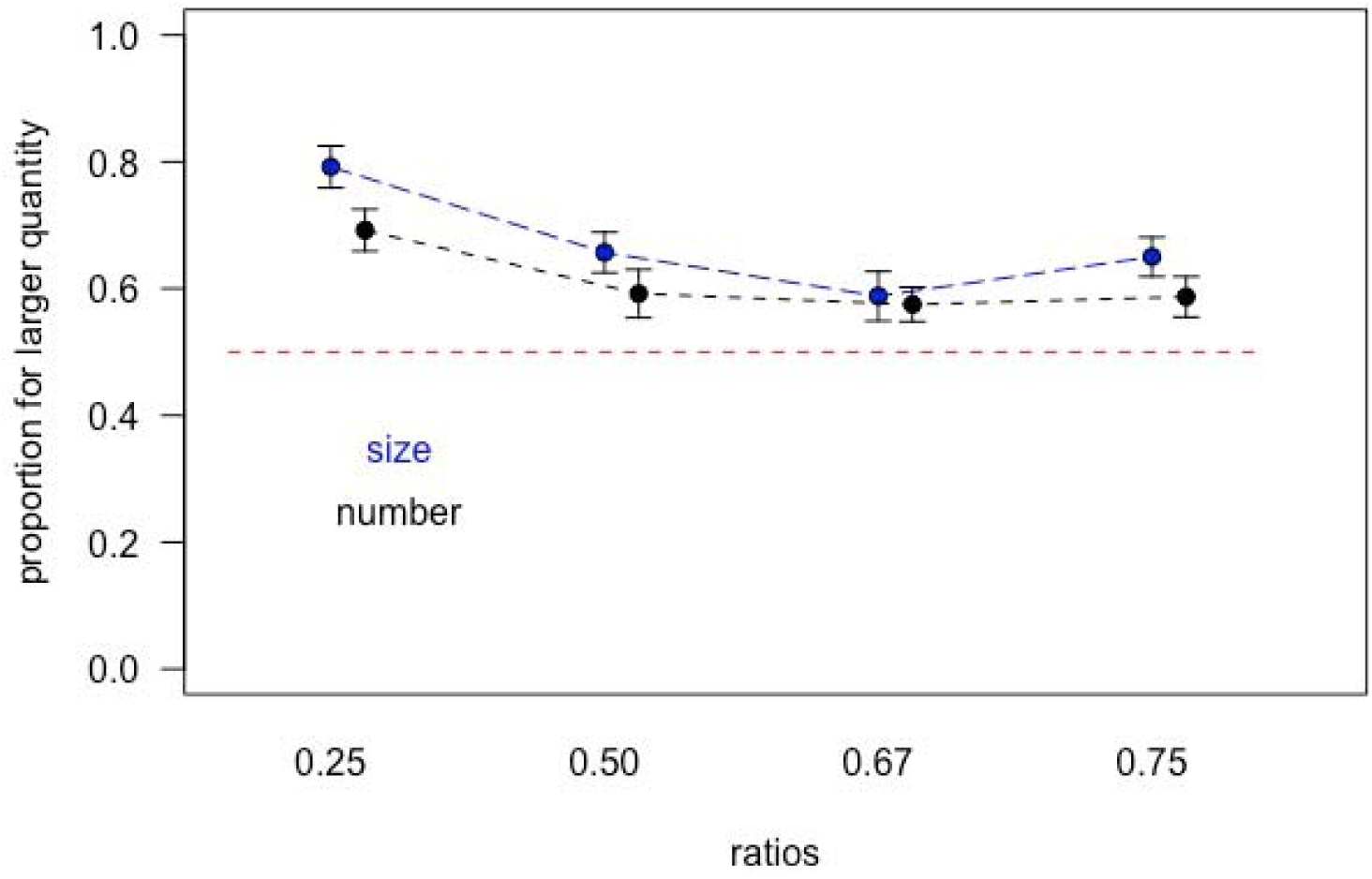
Plot showing the means and se for the index-value (y-axis) for all ratios in both experiments. The red dotted line represents the chance level for choice (index=0.5).

**Fig 3.**
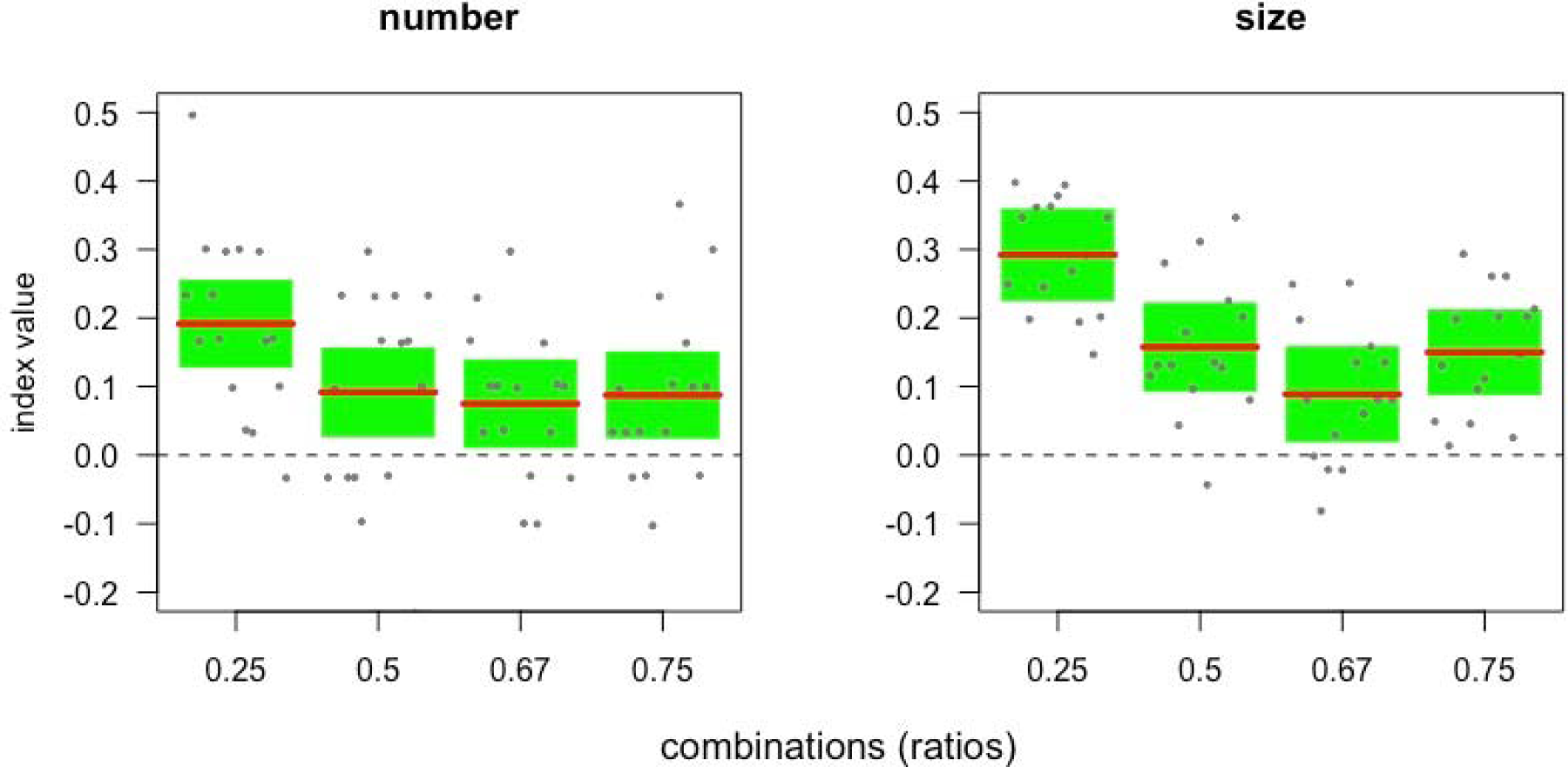
Index expected value and 95% confidence interval bands (green colour) calculated from the mixed models (by bootstrap with 10000 repetitions), for the separated experiments. Dashed lines represent chance value of the index.

Considering the animals tested in both of the experiments (12 subjects overall), we did not find a significant correlation between number and size experiment for the same ratio (Pearson’s product-moment correlation test (lower p-value for 0.25 ratio): t=1.70, df=10, p=0.12; for this ratio effect size was medium-large: r=0.47).

## Discussion

Tortoises showed a remarkable ability to discriminate food items of different numerosity and of different size. Their performance was ratio-dependent and aligns with the ability observed in other groups of vertebrates (Vallortigara, 2014; 2017).

However, among vertebrates, the class of reptiles has remained largely understudied for numerical competence. Evidence of poor performance in lizards as compared to fish (see Introduction) has been hypothesized to be due to some genetic change which took place within ray-finned fish, but after their divergence from the lineage leading to land vertebrates. This promoted the appearance of complex cognitive skills in fish, including numerical abilities (Miletto Petrazzini et al, 2017). Our results with tortoises suggest that this is not the case, since these reptiles show numerical performances comparable to those of fish and amphibians. It seems likely that difficulties of lizards with quantity discrimination may depend on more mundane factors related to motivation, task used or type of reward, thus supporting the first hypothesis formulated for *Podarcis siculus (Miletto Petrazzini et al, 2017)*. Lizards are known to actively prey on live animals in movement (). Therefore, dead *Musca domestica* larvae used in the previous experiment could not have properly simulated a fairly intense food stimulus to motivate the interest of the lizards in discriminating larger numerical quantities.

The procedures used with lizards (Miletto Petrazzini et al, 2017) and, in the present study, with tortoises, did not allow to disentangle the specific role played by strictly numerical aspects of the stimuli and those associated with continuous physical variables. Even with discrete items, the discrimination could have been based on some computation of continuous physical variables that co-vary with numerosity (see Leibovich et al., 2017). It is interesting to note, however, that the tortoise’s performance was different both in the discrete numerical discrimination and the size discrimination (the latter being easier). Moreover, no correlation was observed between number and size experiment for the same ratio. This could be suggestive of different mechanisms involved.

Among the extant reptile orders, most research in cognition has focused on chelonians (Wilkinson and Huber 2012) and has revealed remarkable abilities in spatial cognition (Lopez et al. 2003; Wilkinson et al 2007), visual cognition (Wikinson et al 2013) and acquisition of novel behaviours (Davis and Burghardt 2007). Over the last 225 million years, chelonians seem to have undergone little change, thus representing earlier evolutionary solutions to cognitive problems such as quantity estimation (Wilkinson and Huber 2012).

**Ethics.**

## Data accessibility

The dataset is available from the electronic supplementary material, S3.

## Authors’ contributions

AG,DP,GV. developed the study concept; … AG,DP,GV …. contributed to the study design;. … AG,DP … collected the data; AG,DP,GV…….. analysed the data; … AG,DP,GV…. drafted the manuscript. All authors contributed to the revisions of this manuscript, agreed to be held accountable for this work and approved the final version of the manuscript for publication.

## Competing interests

The authors declare no competing interests.

**Funding.**

## Acknowledgements

We are grateful to “Oasi di San’Alessio” and to Giulio and …. Salamon for hosting the experiment.

We thank Luca Boscolo and Valentina Biglieri for help during the experiment. Thank you to Marco Mangiacotti for his useful advice on the analysis of data.

## Supplementary materials

### Species and study area

Hermann’s tortoise (*Testudo hermanni*) is a small to medium-sized tortoise widespread throughout southern Europe. In Italy, the species occupies a great variety of habitats (Cheylan et al., 2011), both open (coastal areas, dunes, garrigues and bushy glades) and wooded (Mediterranean scrub or mesophilous woods). The diet of this tortoise is essentially composed of plants, in addition to invertebrates, small dead animals and bones (Cheylan, 2001). It mainly consumes annual plants, while avoiding woody, aromatic and resinous ones. It also feeds on fruits of various species, as well as algae, mosses and fungi. Hermann’s tortoise is active all year except during winter months (December to February), the mating season starting from mid-March until the end of the summer, depending on climatic conditions (Calzolai & Chelazzi 1991).

The study was conducted at the “Oasi di Sant’Alessio”, located in Sant’Alessio con Vialone (Lombardy, Northern Italy). Here, tortoises are kept in enclosures at semi-natural conditions at medium-high density and are fed daily with fruit and vegetables. In our experiments we decided to use tomatoes (*Solanum lycopersicum*, San Marzano tomato variety) both because they are one of the tortoises’ most favourite vegetables and because the colour red is known to be significantly attractive to this species (Pellitteri-Rosa et al., 2010). In order to quickly recognize the individuals during experimental trials, we marked them by painting a number on the top carapace of each tortoise. In order to determine sexual adultness, we measured carapace length of all turtles using a digital calliper (accuracy ± 0.1 mm). Sexual dimorphism is noticeable in this species: males are smaller and their plastron is concave, which allows them to mount females during mating (Willemsen and Hailey 2003). In our experiments we selected 16 males and 3 females (carapace length mean ± ES, males: 167.7 ± 4.3 mm, females 200.0 ± 26.2 mm). We marked all tortoises by painting a number on their top carapace to quickly recognize the individuals during experimental trials.

### Experimental set-up

The experimental apparatus consisted in a Y-shaped arena inserted into a wooden rectangular enclosure (120 × 60 × 45 cm). The arena was divided into a tunnel (90 × 28 × 45 cm) that served both as starting zone and approach area to the visual stimuli and a testing compartment (30 × 60 × 45 cm) where food (tomato slices) was placed during trials. The path in the tunnel was inclined at an angle of 25° to allow the tortoises to easily reach the testing compartment. Since all trials were run outside, we set up side walls (45 cm) and an upper shelf above the experimental apparatus in order to discard the effects of sun light, i.e. the formation of shaded and lighted zones, and surrounding environment on behavioural responses.

In order to keep the tortoises in a central position and equidistant from the stimuli before choosing, we set the tunnel width (28 cm) according to the tortoises’ size used in the experiments. Food was presented on two wooden pyramidal base supports (10 × 4 cm) placed on the lower surface into the centre of the testing compartment, both equidistant from the subject’s path of approach.

We filled the testing compartment with the smell of four tomato slices, placed out of the tortoises’ sight, in order to reduce the chance of using olfactory senses to discriminate quantities. Four wooden shelves (10 × 4 cm) containing two tomato slices each were placed on the bottom wall of the testing compartment to allow the smell of the food to
permeate the compartment, as already adopted in other cognitive studies. The tomato slices were changed at the onset of each session.

Stimuli presented on wooden supports were tomato slices differing in number or size according to the experiment. In the number discrimination experiment, two groups of circular tomato slices of equal size (2 cm diameter) were used so that the total amount of food correlated with the number of items (e.g. the ratio between the amount of food in 1 vs. 4 contrast was equal to 0.25). The tortoises were tested with four numerical contrasts: 1 vs. 4, 2 vs. 4, 2 vs. 3, and 3 vs. 4 (respectively: 0.25, 0.50, 0.67, and 0.75 ratios). We arranged the tomato slices in spatially different positions across trials, without adopting any specific pattern, in order to avoid the tortoises from being influenced in their choices by the position of food presented in the previous trials.

In the size discrimination experiment, two items were provided in each trial (1 vs. 1) by presenting one slice placed in the centre of each wooden support during each trial. Differently rectangular sized food items (range from 2 × 1 cm to 2 × 4 cm) were presented and the ratio between the size of the food item within each pair was the same used in the previous experiment: 0.25, 0.50, 0.67, and 0.75 (e.g. in 0.25 ratio, one slice measured 2 cm^2^ and the other 8 cm^2^).

### Experimental Procedure

The procedure consisted of a pre-testing phase followed by a testing phase.

#### Pre-testing phase

The tortoises underwent a 5-day acclimation phase in order to familiarize with the experimental setting and procedure. Each tortoise was transferred into the apparatus every day and was presented with two supports showing 1 vs. 0 tomato slices. The tortoises were individually placed in the waiting area, but could only access the testing zone after the removal of a thin wooden panel. After 5 sec we removed the panel, allowing the tortoises to explore the experimental apparatus for 10 min and eat the tomato slices. On Days 4 and 5, we prevented the tortoises from eating food by removing them as soon as they approached at the distance of about 1 cm from the tomato slices. The choice was defined at the first stimulus approached by the tortoise. The single item position was counterbalanced over trials to inhibit side biases. The tortoises that did not move or moved but did not approach either stimuli within the cut-off time (3 min) were not admitted to the testing phase.

#### Testing phase

We gently placed a tortoise in the starting zone, the stimuli differing either in number or in size of food items located in the testing compartment and the wooden panel inserted to prevent sight of the tortoise. After 5 sec, the panel was removed and the tortoise could access the testing compartment, thus being able to choose between the two stimuli. To exclude any possible learning effect, the tortoises were not allowed to eat the tomatoes, thus allowing the subject’s response to be emitted in the context of a spontaneous choice task. If the tortoise did not make any choice within 3 min, the trial was considered not valid and later repeated.

In both experiments, the subjects were tested on alternate days. In the number experiment, the tortoises underwent a total of 60 trials distributed over 13 days. Numerical discriminations were intermingled across trials within each testing session (each numerical discrimination was presented once in each session, twice in few sessions). In the size experiment, the tortoises underwent 60 trials distributed over 13 days. Size discriminations were intermingled across trials within each testing session (each size discrimination was presented at least once in each session and the four ratios were counterbalanced across sessions).

To conduct the analysis, we used an index of choice as response variable. The index was calculated as the number of choice for larger item / total number of choice. To explore differences between experiments we run a linear mixed-effect model (LME) on the index, while the type of combination (i.e. ratio) and of the experiment (number or size discrimination) were entered as fixed factors, and their interaction was included in the model; subject was entered as random factor and was nested with the type of the experiment. Response variable was inserted in the in the model by subtracting the constant 0.5 (index =0.5), which represents the chance value for the choice, to allow an easier interpretation of the results. Bootstrap estimate of confidence intervals (parametric boostrapping n=10000) of models were obtained by R package “bootpredictlme4” (Duursma 2017) and visualized within R package “visreg” (Breheny and Burchett 2017).

For both experiment normality of response was checked with Shapiro-Wilk test.

Pattern analysis was performed considering the combination as numerical predictor in a linear mixed model with the same structure of the previous model (Li and Baron 2012).

We used R (R core team 2018) and package “lme4” (Bates et al. 2015))to perform the statistical analysis.

In Experiment 2, one subject ceased to respond after the 11^th^ trial in the 0.25 ratio and after the 8^th^ trial in all the other three ratios (0.50, 0.67, 0.75). Its performance was considered only up to this point.

## References

Agrillo, C., Piffer, L., Bisazza, A., and Butterworth, B. (2012). Evidence for two numerical systems that are similar in humans and guppies. PLoS ONE 7:e31923. doi:10.1371/journal.pone.0031923

Cantlon, J.F., Brannon, E.M. (2007). Basic Math in Monkeys and College Students. PLoS Biology, 5: e328.

Davis, K. M., & Burghardt, G. M. (2007). Training and longterm memory of a novel food 443 acquisition task in a turtle (Pseudemys nelsoni). Behavioural Processes, 27, 225–230.

Ditz H.M., Nieder A. (2016). Numerosity representations in crows obey the Weber-Fechner Law. Proceedings of the Royal Society B, 283: 20160083.

Ferrigno, S., Cantlon, J.F., 2017. Evolutionary Constraints on the Emergence of Human Mathematical Concepts. In: Kaas, J (ed.), Evolution of Nervous Systems 2e. vol. 3, pp. 511–521. Oxford: Elsevier.

Leibovich, T., Katzin, N., Harel, M., and Henik, A. (2017). From ‘sense of number’ to ‘sense of magnitude’ – the role of continuous magnitudes in numerical cognition. Behav. Brain Sci., 40: e164.

Lopez, J. C., Vargas, J. P., G.mez, Y., & Salas, C. (2003). Spatialand nonspatial learning in turtles: Th e role of the medial cortex. Behavioural Brain Research, 143, 109–120.

Miletto Petrazzini ME, Bertolucci C and Foà A (2018) Quantity Discrimination in Trained Lizards (*Podarcis sicula*). Front. Psychol. 9:274. doi: 10.3389/fpsyg.2018.00274

Miletto Petrazzini ME, Fraccaroli I, Gariboldi F, Agrillo C, Bisazza A, Bertolucci C, Foaà A. (2017). Quantitative abilities in a reptile (*Podarcis sicula*). Biol. Lett. 13: 20160899. http://dx.doi.org/10.1098/rsbl.2016.0899

Pepperberg IM. 2006 Grey parrot numerical competence: A review. Animal Cognition 9, 377–391. (doi:10.1007/s10071-006-0034-7)

Potrich, D., Sovrano, V.A., Stancher, G., Vallortigara, G. (2015)._Quantity discrimination by zebrafish (*Danio rerio*). Journal of Comparative Psychology, 29: 388–393.

Rogers, L.J.,. Vallortigara, G., Andrew, R.J. (2013). Divided Brains. The Biology and Behaviour of Brain Asymmetries. Cambridge University Press, New York.

Rugani, R., Fontanari, L., Simoni, E., Regolin, L., Vallortigara, G. (2009). Arithmetic in newborn chicks. Proceedings of the Royal Society of London B, 276: 2451–2460.

Stancher G, Rugani R, Regolin L, Vallortigara G. 2015 Numerical discrimination by frogs (*Bombina orientalis*). Animal Cognition 18, 219–229. (doi:10.1007/s10071-014-0791-7)

Uller C, Jaeger R, Guidry G, Martin C. 2003 Salamanders (*Plethodon cinereus*) go for more: Rudiments of number in an amphibian. Animal Cognition 6, 105–112. (doi:10.1007/s10071-003-0167-x)

Utrata E, Virányi Z, Range F. 2012 Quantity discrimination in wolves (*Canis lupus*). Frontiers in Psychology 3, 1–9. (doi:10.3389/fpsyg.2012.00505)

Vonk J, Beran MJ. 2012 Bears ‘count’ too: Quantity estimation and comparison in black bears, Ursus americanus. Animal Behaviour (doi:10.1016/j.anbehav.2012.05.001)

Vallortigara, G. (2014). Foundations of Number and Space Representations in Non-Human Species. In “Evolutionary Origins and Early Development of Number Processing”, pp. 35–66 (Eds., D.C. Geary, D.B. Bearch, K. Mann Koepke), Elsevier, New York.

Vallortigara, G. (2017). An animal’s sense of number. In “The nature and Development of Mathematics. Cross Disciplinary Perspective on Cognition, Learning and Culture” (Adams, J.W., Barmby P., Mesoudi, A., eds.), pp. 43–65, Routledge, New York.

Wilkinson, A., Chan, H. M., & Hall, G. (2007). A study of spatial learning and memory in the tortoise (Geochelone carbonaria). Journal of Comparative Psychology, 121, 412–418.

Wilkinson, A. & Huber, L. 2012. Cold-blooded cognition: reptilian cognitive abilities. In: The Oxford Handbook of Com-parative Evolutionary Psychology (Vonk, J. & Shackelford, T. K., eds). Oxford University Press, Oxford, pp. 129–143.

## References

Douglas Bates, Martin Maechler, Ben Bolker, Steve Walker (2015). Fitting Linear Mixed-Effects Models Using lme4. Journal of Statistical Software, 67(1), 1–48. doi:10.18637/jss.v067.i01.

Breheny P, and Burchett W. 2017. visreg: Visualization of Regression Models. R package version 2.4-1. https://CRAN.R-project.org/package=visreg

Duursma R. 2017. bootpredictlme4 GitHub repository, https://github.com/remkoduursma

Li, Y., and Baron, J. (2012). Behavioral Research Data Analysis with R. New York, NY: Springer.

R Core Team (2018). R: A language and environment for statistical computing. R Foundation for Statistical Computing, Vienna, Austria. URL https://www.R-project.org/

